# Two potent, functionally redundant MazF toxins inactivate the 16S rRNA anti-Shine-Dalgarno sequence and are differentially expressed in *Mycobacterium tuberculosis*

**DOI:** 10.64898/2026.10.23.761202

**Authors:** Timothy W. Sherrier, Eduardo A. Troian, Unnati Chauhan, Tatsuki Miyamoto, Nancy Woychik

## Abstract

*Mycobacterium tuberculosis* (Mtb) remains a major global health threat, sustained by lengthy, multidrug treatment regimens that promote poor compliance and antibiotic resistance. The pathogen’s ability to evade clearance is facilitated in part by a transition to a nonreplicating, persistent state, a process thought to involve the abundant type II toxin-antitoxin (TA) systems. The Mtb chromosome encodes 11 MazEF TA systems whose MazF toxins are single-strand, sequence-specific endoribonucleases yet the precise RNA targets and physiological consequences of most remain unknown. Here we applied a specialized 5’-end RNA-seq approach to pinpoint the Mtb MazF-mt1 toxin recognition sequence and RNA target. MazF-mt1 specifically cleaves at U↓CCUU within the anti-Shine-Dalgarno (aSD) sequence, releasing it from the 3’ end of the 16S rRNA. Notably, the recently described MazF-mt11 toxin also targets the aSD for removal, cleaving at a different sequence 4 nts upstream of the MazF-mt1 site. Consistent with the essential role of the aSD in translation, both toxins rapidly inhibit new protein synthesis. This functional redundancy prompted us to mine Mtb transcriptomic datasets to compare how the two toxins respond to infection-relevant stresses. MazF-mt1 and MazF-mt11 are differentially regulated following exposure to diverse antibiotic and host-imposed stresses, with distinct temporal expression patterns. Thus, Mtb may coordinate transcription of these functionally redundant toxins to more precisely control the establishment and maintenance of nonreplicating persistence, thereby confounding pathogen clearance.

**Importance:** The ability of *M. tuberculosis* to resist immune-mediated clearance and antibiotic treatment is central to its success as a pathogen. Here, we identify the target of the endoribonuclease toxin MazF-mt1 as the anti-Shine-Dalgarno sequence near the 3’ end of 16S rRNA, cleavage of which causes rapid growth arrest and translation inhibition. Base-pairing between anti-Shine-Dalgarno/Shine-Dalgarno sequences is required for efficient ribosome recruitment and translation of most bacterial mRNAs. Because we previously reported that the MazF-mt11 toxin also cleaves the 16S rRNA 3’ end to remove the anti-Shine-Dalgarno sequence, MazF-mt1 expression phenocopies MazF-mt11. We further show that this toxin pair responds in contrasting ways to several first- and second-line antibiotics and a wide range of environmental stress conditions. Thus, *M. tuberculosis* encodes functionally redundant, ribosome-targeting toxins that are differentially deployed under stress, illuminating a potential mechanism to rapidly suppress translation and promote entry into a drug-tolerant, nonreplicating persistent state during infection.

## Introduction

*M. tuberculosis* (Mtb), the causative agent of tuberculosis, remains a major global public health concern and the leading cause of death from a single infectious agent, accounting for an estimated 1.2 million deaths and 10.7 million new cases in 2024 (1). During infection, Mtb resides primarily within host macrophages, where encounters hypoxia, nutrient limitation, acidic pH, and other immune-imposed stresses (reviewed in (2)). In response to these conditions, Mtb can transition from active replication to a nonreplicating persistent state characterized by extensive metabolic remodeling and reduced antibiotic susceptibility. The ability to enter this state is thought to contribute to the prolonged treatment regimens required for tuberculosis, underscoring the importance of defining the mechanisms that enable Mtb to respond and adapt to immune-imposed stresses.

There is increasing evidence linking the abundant toxin-antitoxin (TA) systems encoded in Mtb (over 70, (3)) to this pathogen’s ability to adapt to and persist under these stresses (reviewed in (4–7)). TA systems are typically small, two-gene operons encoding a stable toxin that acts on an intracellular target and a labile antitoxin that binds to and inhibits toxin activity. Under stress conditions, the antitoxin is thought to be selectively degraded, disrupting the balance between toxin and antitoxin and enabling the toxin to act on its cellular target (reviewed in (8)).

Relative to most bacteria, which encode only a few TA systems, Mtb has markedly expanded TA families, including 50 members of the VapBC TA systems and 11 MazEF systems. To date, Mtb MazF toxins function as sequence-specific endoribonucleases that selectively cleave single-stranded regions within tRNA or rRNA targets; some also cleave mRNAs when ectopically expressed in *Escherichia coli* (Table 1). Through this highly targeted RNA cleavage activity, MazF toxins can reprogram cellular physiology. MazF-mt9 cleavage of tRNA^LysUUU^ initiates interrelated metabolic changes that are essential for Mtb survival following infection (9).

**Table 1.** Summary of *M. tuberculosis* MazF toxins.

| Woychik/Inouye name | Tuberculist/ Other name | Antitoxin MazE locus | Toxin MazF locus | Publications |
| --- | --- | --- | --- | --- |
| MazF-mt1 | MazF9 | Rv2801A | Rv2801c | *This work*, Sala et al. 2014 (3), Tiwari et al. 2015 (10), Zhu et al. 2006 (13), Zhu et al. 2010(48) |
| MazF-mt2 | MazF1 | Rv0456B | Rv0456A | Sala et al. 2014 (3) |
| MazF-mt3 | MazF6 | Rv1991A | Rv1991c | Chattopadhyay et al. 2022 (49), Sala et al. 2014 (3), Schifano et al. 2013 (50), Schifano et al. 2014(21), Tiwari et al. 2015(10), Zhao and Zhang 2008 (51), Zhu et al. 2008 (52), Zhu et al. 2010 (48) |
| MazF-mt4 | MazF2 | Rv0660c | Rv0659c | Sala et al. 2014 (3) |
| MazF-mt5 | MazF5 | Rv1943c | Rv1942c | Sala et al. 2014 (3) |
| MazF-mt6 | MazF3 | Rv1103c | Rv1102c | Hoffer et al. 2017 (53), Sala et al. 2014 (3), Schifano and Woychik 2014 (54), Schifano et al. 2013(50), Tiwari et al. 2015(10), Zhu et al. 2006 (13) |
| MazF-mt7 | MazF4 | Rv1494 | Rv1495 | Huang et al. 2010(55), Sala et al. 2014(3), Zhu et al. 2008(52) |
| MazF-mt8 | MazF8 | Rv2274A | Rv2274c | Sala et al. 2014 (3) |
| MazF-mt9 | MazF7 | Rv2063 | Rv2063A | Barth et al. 2019 (44), Sala et al. 2014 (3), Schifano and Woychik 2017 (56), Schifano et al. 2016(20) |
| MazF-mt10 | MazF10 | Rv0298 | Rv0299 | Kim et al. 2016 (57), Sala et al. 2014 (3) |
| MazF-mt11 | mt-PemK | 3467307-3467612* | Rv3098A | Sherrier et al. 2025 (22), Chi et al. 2018 (58) |

\*Genome position in *M. tuberculosis* H37Rv

Deletion studies in guinea pigs further revealed that multiple MazF toxins act cooperatively to enable Mtb to adapt to stress and contribute to drug tolerance and virulence (10). These findings indicate that MazF toxins do more than simply arrest growth; they can inhibit or remodel translation in ways that that support adaptation to stress. Yet, the RNA target specificities and roles of the majority of Mtb MazF toxins remain unknown.

Over 20 years ago, MazF toxins were widely considered “mRNA interferases,” a term coined by Masayori Inouye. This view arose from studies showing that the *E. coli* MazF toxin cleaves mRNAs immediately 5’ of ACA sequences, an activity predicted to result in widespread cleavage across the transcriptome because nearly all *E. coli* mRNAs contain this motif (11, 12). Consistent with this view, primer-extension analysis of RNA from *E. coli* cells expressing MazF-mt1, combined with in vitro cleavage of *M. smegmatis* RNA and synthetic RNA substrates by recombinant toxin, led the Inouye group to conclude that MazF-mt1 cleaves mRNAs at U↓AC sequences (13). However, our laboratory subsequently found that recombinant toxin can exhibit promiscuous cleavage in vitro and that expression in the native bacterial host is required to precisely define cleavage specificity (14).

Here we show that the physiological target of MazF-mt1 is the U↓CCUU sequence near the 3’ end of 16S rRNA. Cleavage at this site disrupts the anti-Shine-Dalgarno sequence (aSD), resulting in widespread inhibition of translation. Notably, this target and mechanism closely parallel what we previously reported for MazF-mt11, which cleaves a neighboring region in the 16S rRNA and similarly alters translation. The mechanistic similarity between these two toxins prompted us to investigate their role during the stress response. Mining publicly available transcriptomic datasets revealed that *mazF-mt1* and *mazF-mt11* are differentially regulated in response to multiple antibiotics and host-relevant environmental stresses, suggesting that Mtb may coordinate expression of these toxins to precisely control the establishment and maintenance of nonreplicating persistence.

## Results

### Expression of MazF-mt1 leads to growth arrest in *E. coli*, *Mycobacterium smegmatis*, and Mtb

MazF-mt1 (Rv2801c) is one of 11 members of the MazEF family in Mtb (Table 1). Characteristic of all TA systems, the toxin and antitoxin transcripts are in an operon, sometimes with other genes. MazEF1 comprises a two-gene operon whose genes either overlap, as they do here by 17 bp, or are separated by a short distance (Fig. 1A). Among its Mtb paralogs, the MazF-mt1 protein sequence exhibits highest sequence similarity to MazF-mt2 (57% identity and 14% similarity, 71% overall similarity; Fig. 1B). MazF-mt1 is also more closely related to the *E. coli* MazF (33% identity, 14% similarity) than to most other Mtb MazFs (Fig. 1B,C) with high structural similarity by X-ray crystallography (RMSD=1.63Å indicating the two proteins adopt closely similar overall folds); Fig. 1D). Consistent with this conservation, MazF-mt1 retains the residue corresponding to E24, which is important for substrate recognition by *E. coli* MazF (15), as well as identical or conserved counterparts corresponding to the catalytic R29 and T52 of *E. coli* MazF (15) (Fig. 1C).

**Figure 1.**
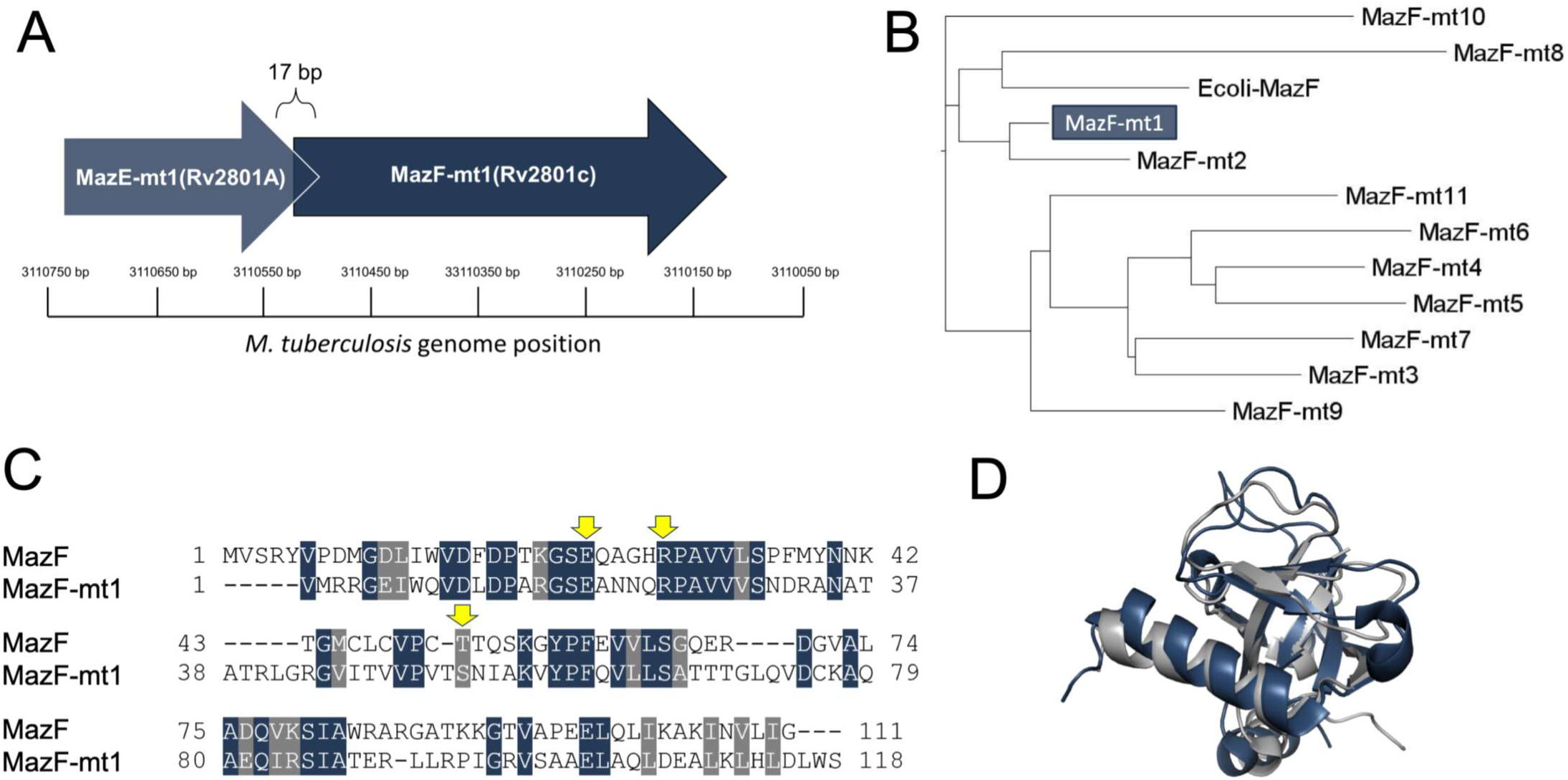
MazF-mt1 exhibits sequence and structural similarity to *E. coli* MazF. A, Genome organization of genes encoding MazEF-mt1 highlighting the 17 bp out of frame overlap. B, Phylogenetic tree with *E. coli* MazF and the 11 Mtb MazFs was generated first using IQ tree 3.0.1 (59) maximum likelihood method to calculate sequence relationships, followed by visualization with Figtree 1.4.4 https://tree.bio.ed.ac.uk/software/figtree/. C, Clustal Omega (60) alignment of MazF-mt1 to *E. coli* MazF. Blue, identical; light grey, similar amino acids (BLOSUM62 ≥ 1). Arrows denote conserved residues E24, R29, and T52 important for *E. coli* MazF activity. D, High resolution Xray structures of *E. coli* MazF (PDB 1UB4) aligned with MazF-mt1 (PDB 6KYS) using Pymol (The PyMOL Molecular Graphics System, Version 3.0 Schrödinger, LLC).

Since a characteristic phenotype of MazF toxin expression is growth arrest, we monitored the effect on MazF-mt1 on growth in *E. coli, Mycobacterium smegmatis*, and Mtb. We expressed MazF-mt1 in *E. coli* using the arabinose-inducible pBAD33 plasmid. Following toxin induction, growth slowed relative to the empty vector control, leading to nearly complete and sustained growth arrest after 30 min of toxin expression (Fig. 2A).

**Figure 2.**
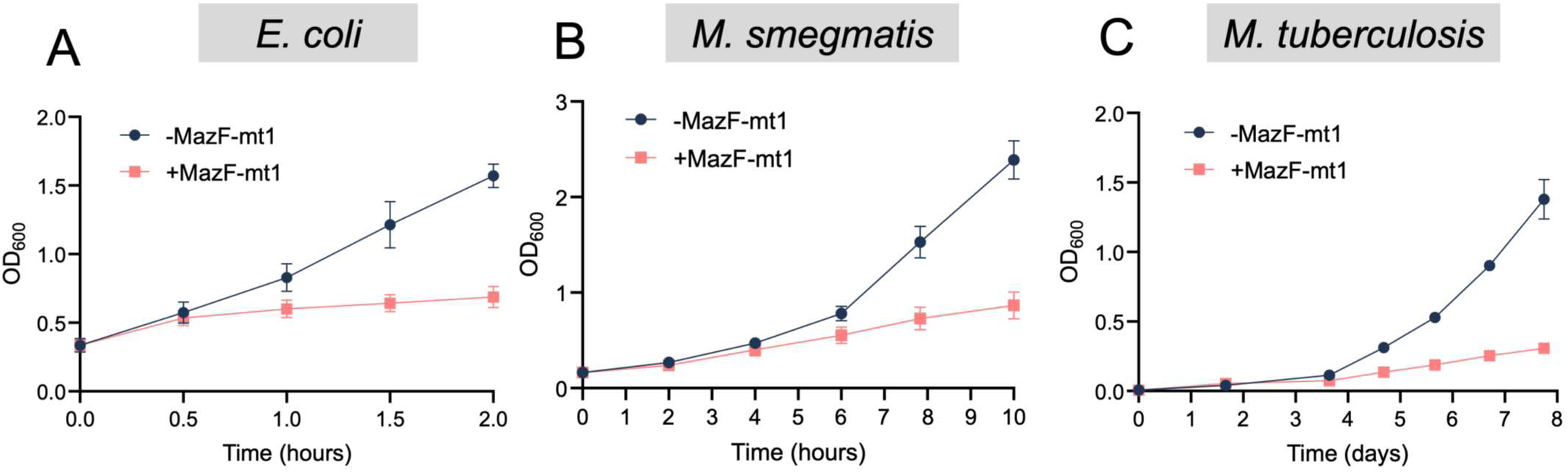
Expression of MazF-mt1 leads to growth arrest in *E. coli*, *M. smegmatis*, and Mtb. A, *E. coli* cultures containing pBAD33-*mazF-mt1* were grown to OD_600_ of 0.3 in M9 + glycerol and induced with 0.2% arabinose. Error bars represent standard deviation from the average of four biological replicates. B, *M. smegmatis* cultures containing the pMC1s-*mazF-mt1* plasmid were grown overnight to an OD_600_ of 0.1, split in half and grown with (+*MazF-mt1*) or without (-*MazF-mt1*) ATc inducer. Error bars represent standard deviation from the average of three biological replicates. C, Individual Mtb colonies containing the pMC1s-*MazF-mt1* plasmid were first resuspended in 1 ml of 7H9 and used to inoculate two cultures with (+*MazF-mt1*) or without (-*MazF-mt1*) ATc. Error bars represent standard deviation from the average of three biological replicates.

MazF-mt1 was expressed in *Mycobacterium smegmatis*—which only has a single MazF toxin that targets tRNA^Lys^ (16)—using the anhydrotetracycline (ATc) inducible pMC1s integrating plasmid engineered for low-level expression in mycobacteria (16). Toxin-expressing cells exhibited a gradual slowing of growth beginning 4h post-induction through 10h (Fig. 2B). Finally, expression of MazF-mt1 in Mtb using the pMC1s plasmid also slowed growth beginning approximately 4 days post-induction persisting through day 8 (Fig. 2C) Thus, MazF-mt1 expression inhibits cell growth, consistent with other Mtb MazF toxins characterized to date.

### MazF-mt1 cleaves a five-nucleotide sequence motif in *E. coli*

As noted above, none of the Mtb MazF toxins subjected to rigorous characterization were found to target mRNAs. Thus, the assignment of MazF-mt1 as a U↓AC-specific mRNA interferase in a 2006 report from the Inouye group based on detection of common cleavage sites within a single *E. coli* transcript (the 906 bp *era* transcript) and in short synthetic RNAs (13), is likely incorrect. Instead, we applied our specialized 5’ RNA-seq method to identify the type of RNA MazF-mt1 targets as well as the consensus sequence required for RNA cleavage. We previously discovered that application of this genome-scale method to RNA derived from cells expressing a MazF toxin in *E. coli* or *Mycobacterium smegmatis* can be useful for identification of the cleavage consensus sequence (17–22). Therefore, we used MORE (<u>m</u>apping by <u>o</u>verexpression of an <u>R</u>Nase in *<u>E</u>. coli*) RNA-seq developed in our laboratory (21), to help identify the MazF-mt1 RNA recognition sequence and the precise cleavage position within this consensus site.

MazF-mt1 was expressed from pBAD33 in an *E. coli* strain with deletions of its six endoribonuclease TA systems, including the sole MazEF operon. RNA was harvested from toxin-expressing and control cells 2 hrs post induction and subjected to MORE RNA-seq (Supplemental Table 1). Fig. 3A shows the top 20 *E. coli* RNA targets sorted by descending fold change between toxin-expressing and control cells. The 5 nt weblogo ^5’^U(C/A)CUU^3’^ derived from all targets meeting a ≥60-fold enrichment cutoff revealed that MazF-mt1 recognizes and cleaves RNA after the 5’-U within two closely related consensus sequences, U↓CCUU and U↓ACUU (Fig. 3B). This 5 nt sequence derived from the entire *E. coli* transcriptome is a more accurate representation of MazF-mt1 RNA target sites compared to the previously published 3 nt U↓AC consensus derived from common cleavage sequences identified in *era* mRNA (13).

**Figure 3.**
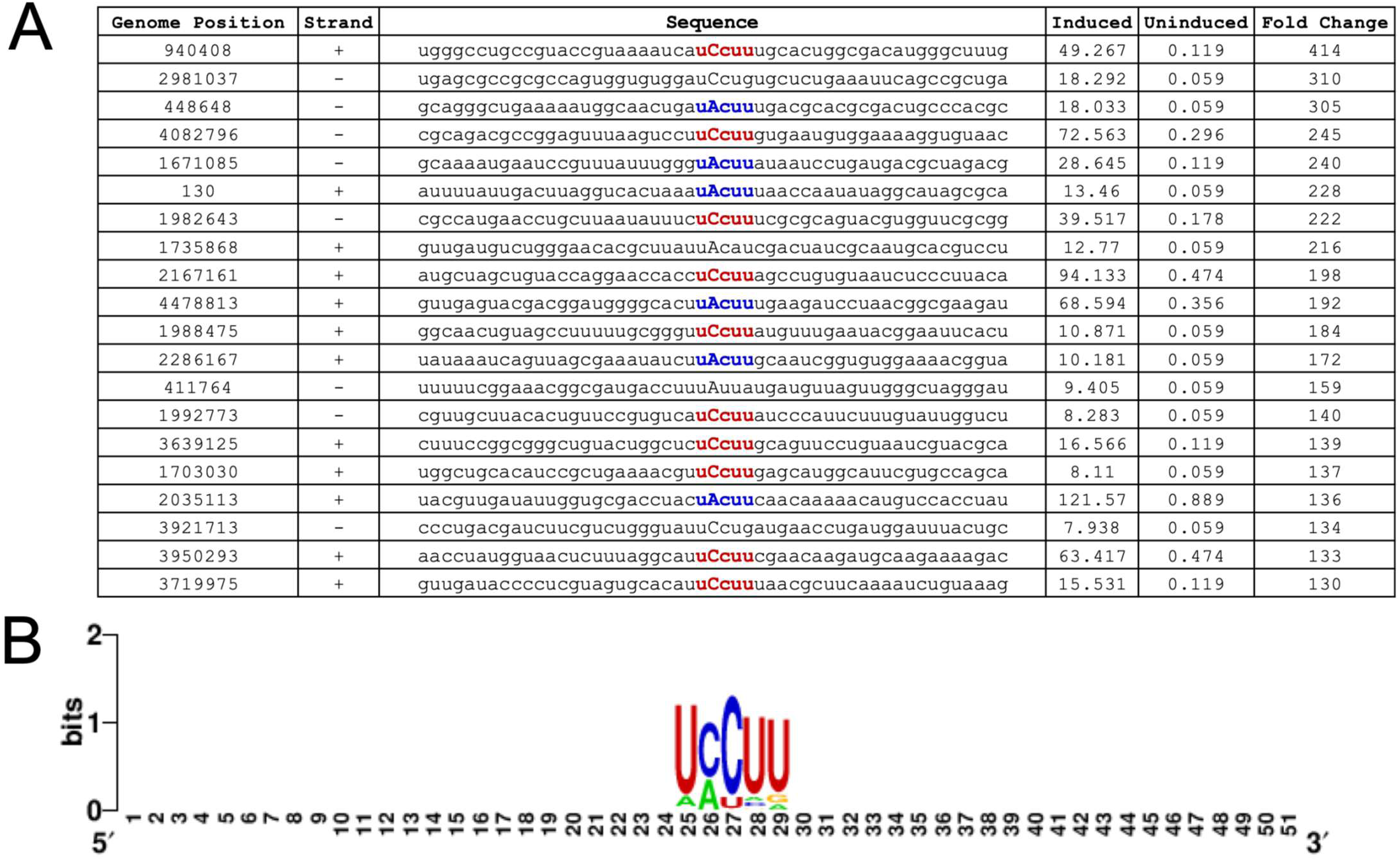
MazF-mt1 cleaves RNA at U↓ACUU and U↓CCUU in *E. coli*. A, MORE RNA-seq (21) identified two distinct 5 nt RNA recognition sequences U↓ACUU (blue) or U↓CCUU (red) from RNA harvested from *E. coli* cells after 2 hrs with or without MazF-mt1 induction. RNA cleavage occurs 5’ of the nts shown in capital letters B, WebLogo (61) derived from all *E. coli* MazF-mt1 cleavage sites enriched 60-fold or more in the MORE RNA-seq dataset.

The degeneracy of the cleavage recognition sequence at the second nt of the U↓(A/C)CUU consensus sequence is curious and atypical. Although MazF endoribonuclease toxins exhibit a strong preference for cleavage at their consensus site, they will cleave at similar sequences or shorter sites when their preferred sequence is limiting, rare, or completely absent. Thus, the observed degeneracy in the *E. coli* host might be due to a paucity of available consensus sequences because of the lower GC content of its genome: 51% versus 67% in Mtb. Indeed, since we observed a preference for a C at the second position—U↓CCUU is represented twice as often as U↓ACUU—our data suggest that U↓CCUU is the MazF-mt1 cleavage site in Mtb.

### MazF-mt1 cleaves 16S rRNA at a single U↓CCUU site within the aSD sequence in *M. smegmatis* and Mtb

To resolve the U↓(A/C)CUU consensus discrepancy, we first expressed MazF-mt1 in *M. smegmatis*; a mycobacterial host whose 67% GC content is nearly identical to that of Mtb. We harvested RNA from cells with or without MazF-mt1 expression via pMC1s as in Fig. 2 and performed 5’-OH RNA-seq to identify the toxin cleavage site and consensus sequence (Supplemental Table 2). Unlike in the *E. coli* host, we identified a single U↓CCUU MazF-mt1 cut site near the 3’ end of the 16S rRNA, with no U↓ACUU cleavage events detected. The toxin removes the terminal 7 nts of the 16S rRNA 3’ end, cleaving within the functionally significant aSD sequence (CCUCCU, Fig. 4A). The 5’-OH RNA-seq data from Mtb with or without toxin expression via pMC1s matched those from *M. smegmatis* (Supplemental Table 3): MazF-mt1 only cleaves a single U↓CCUU site in the 16S rRNA and disrupts the aSD sequence (Fig. 4C). The aSD sequence base-pairs with the Shine-Dalgarno (SD) sequence in mRNAs to align the ribosome with the start codon and enable efficient translation initiation. Cleavage at the U↓CCUU site removes the latter half of the aSD sequence, i.e., it cleaves in the middle of the 6 nt CCU↓CCU aSD (Fig. 4A,C), likely compromising protein synthesis from leadered transcripts carrying an SD sequence.

**Figure 4.**
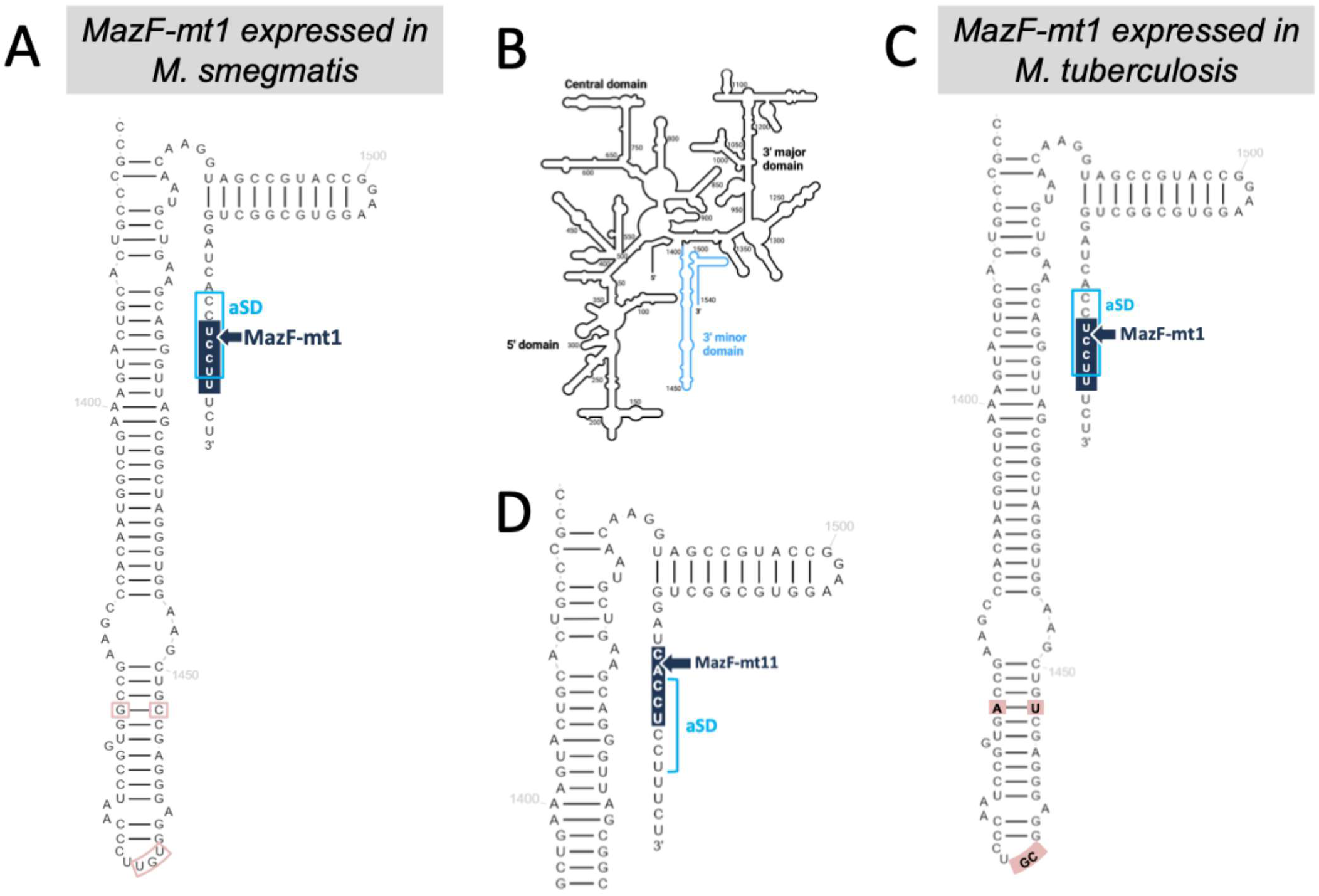
The sole MazF-mt1 target in mycobacteria is 16S rRNA at U↓CCUU near its 3’ end. Detailed sequences and secondary structures of the *M. smegmatis* (A) and Mtb (C) 16S rRNA 3’ minor domain comprising helices 44 and 45 and the single-stranded 3’ end flanking a general schematic of the entire bacterial 16S rRNA from Biorender highlighting the 3’ minor domain (B, light blue). Their slight sequence differences indicated by open (A, *M. smegmatis*) or filled (C, Mtb) salmon boxes. 5’-OH RNA-seq revealed a single MazF-mt1 recognition sequence (dark blue box) whose position of cleavage (arrow) is identical in both 16S rRNAs, as are the CCUCCU aSD sequences (light blue open box). (D) The MazF-mt11 cleavage site at the 3’ end of 16S rRNA is distinct from MazF-mt1.

### Truncation of the 16S rRNA aSD by MazF-mt1 markedly impacts translation

To assess the impact of toxin-mediated scission of the 3’ terminal 7 nt on protein synthesis, we used a click chemistry based metabolic labeling approach to monitor de novo protein synthesis in *M. smegmatis* cells with or without MazF-mt1 expression. Newly synthesized proteins were labeled by incorporation of the azide containing methionine analog azidohomoalanine (AHA) and visualized by fluorescent coupling to the alkyne tetramethylrhodamine (TAMRA). Disruption of the aSD sequence severely impaired new protein synthesis at both 5 and 24 hrs after toxin induction (Fig. 5A), with synthesis reduced to 6% of uninduced levels after 5 hrs and 24% after 24 hrs (Fig. 5B).

**Figure 5.**
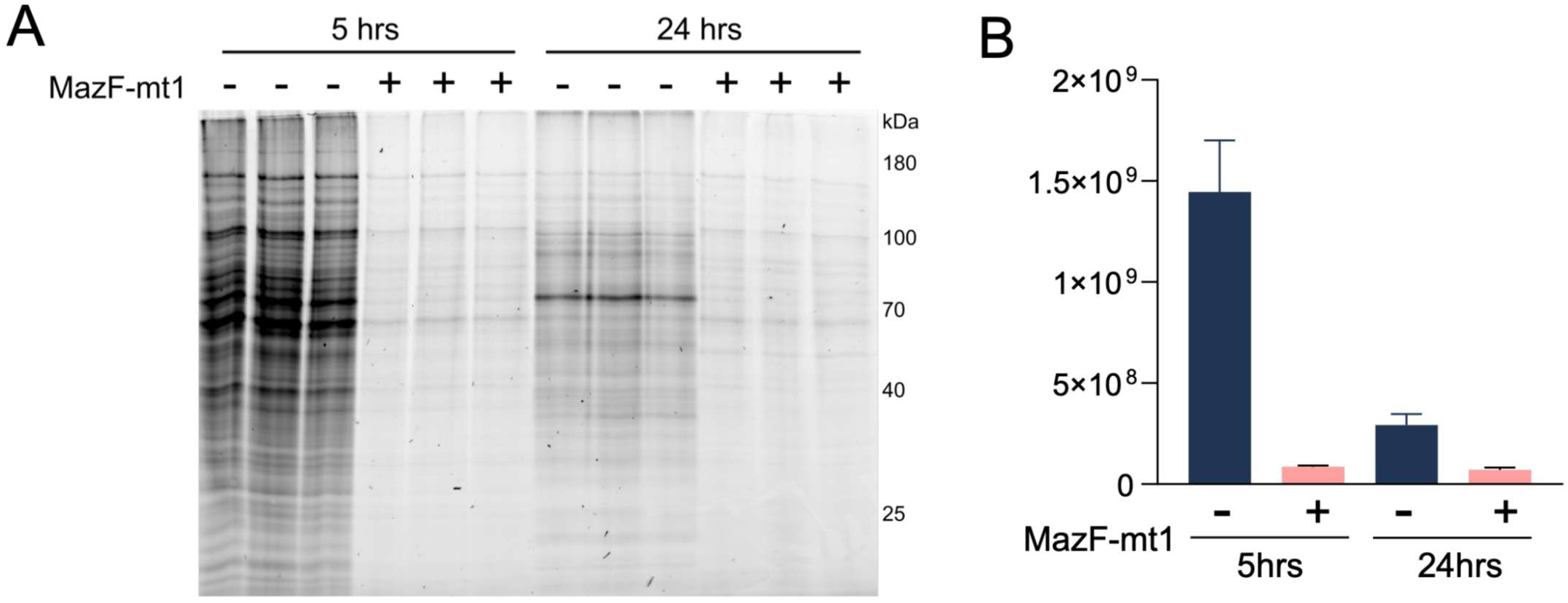
MazF-mt1 expression leads to nearly complete inhibition of translation. A, MazF-mt1 was expressed in *M. smegmatis* for 5 or 24 hrs and followed by labeling with the methionine mimetic azidohomoalanine (AHA) for 3 hrs and visualized upon alkyne-TAMRA conjugation. Each lane was normalized to 10 μg of total protein. B, Whole lane TAMRA fluorescence signals were quantified for each of the three replicates shown; bars represent standard deviation.

### MazF-mt1 and MazF-mt11 are differentially expressed under varying stress conditions

We recently reported that the Mtb MazF-mt11 toxin exclusively targets the 3’ end of 16S rRNA at the C↓ACCU sequence, removing the terminal 11 nt, which include the entire CCUCCU aSD sequence (Fig. 4D) (22). Given that both MazF-mt1 and MazF-mt11 impair aSD function and inhibit translation, we sought to understand why Mtb would retain two TA systems with seemingly redundant functions. To address this, we mined publicly available Mtb H37Rv transcriptomic datasets to assess toxin regulation under stress (23, 24) and antibiotic exposure (25–27). Across these datasets, both toxins showed strong differential responses to several common first-and second-line Mtb antibiotics (Fig. 6A-D). *mazF-mt1* transcript levels increased in response to bedaquiline, with highest transcript abundance after 4 and 24 hrs (Fig. 6D) and prolonged linezolid treatment (72 hrs, Fig. 6B), whereas *mazF-mt11* was upregulated in response to pretomanid for 24 and 72 hrs (Fig. 6C) and shorter linezolid exposure (4 and 24 hr, Fig. 6B). Both toxin transcripts were upregulated in response to isoniazid exposure for 72 hrs, with *mazF-mt11* showing a stronger relative response at 24 and 72 hrs (Fig. 6A).

**Figure 6.**
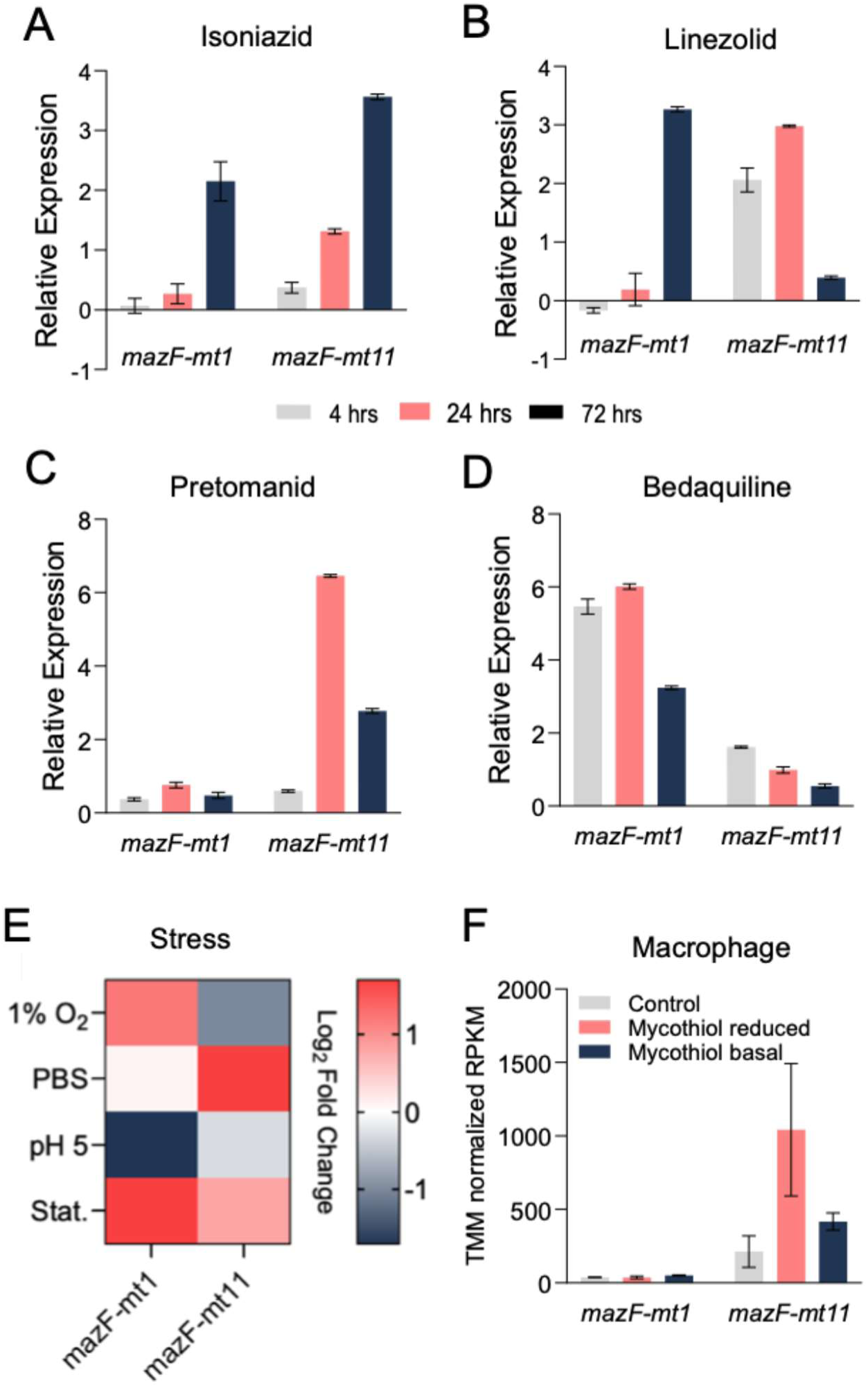
MazF-mt1 and MazF-mt11 are differentially expressed during infection-related stress conditions. A-D, and F, Bars show the difference in TMM-normalized RPKM between treatment and wild-type controls; a value of 0 indicates no difference. Error bars represent the standard deviation of three replicates. A-D depicts the transcriptional responses of each toxin to the indicated antibiotics (25–27). E, Heat map of the transcriptional responses of each toxin to the indicated stress conditions ((24), Stat.=stationary phase). Effect size and significance cutoff is |log₂FC| ≥ 1 and adjusted *p* ≤ 0.05. F, Transcriptional responses following infection of THP-1 cells with Mtb H37Rv, sorted by mycothiol redox state (23, 25).

In addition to these distinct expression patterns during antibiotic stress, the two toxins responded differently to environmental stresses relevant to Mtb infection ((24), Fig. 6E). Using an effect-size threshold of |log₂FC| ≥ 1 and an adjusted *p* ≤ 0.05 significance cutoff, *mazF-mt1* was upregulated during stationary phase and hypoxia (SP and 1% O_2_, Fig. 6E) and downregulated at low pH (pH 5, Fig. 6E), while *mazF-mt11* transcript levels increased under nutrient starvation (PBS, Fig. 6E). Finally, only *mazF-mt11* transcription was upregulated in the THP-1 macrophage infection model (Fig. 6F). Here, THP-1 macrophages were separated into two populations, reduced and basal, gated by redox potential using a redox-sensitive fluorescent probe (23, 25). Although the Singh group had previously separated THP-1 macrophages into populations enriched for reduced, basal, or oxidized mycothiol redox potential (28), they performed RNA-seq only on the reduced and basal populations (23). The basal population reflects a baseline/intermediate redox state, whereas the reduced population represents an adaptive, drug-tolerant redox state. Only *mazF-mt11* expression increased modestly in both intracellular populations relative to the in vitro control (Mtb grown in 7H9), with higher expression observed in the reduced redox population, thought to be more resistant to antibiotic treatment.

## Discussion

Here, using a combination of MORE RNA-seq and 5’-OH RNA-seq, we establish the true target of MazF-mt1 in Mtb as the 3’ end of 16S rRNA, with cleavage at the U↓CCUU recognition sequence. The exquisite specificity of MazF-mt1 is striking. The Mtb transcriptome contains 796 genes whose transcripts carry at least one UCCUU consensus sequence. Yet this MazF-mt1 exclusively recognizes and cleaves only the U↓CCUU site at the 3’ end of the 16S rRNA. This specificity indicates that additional determinants govern cleavage site recognition in vivo. We previously demonstrated that the MazF-mt9 toxin recognizes and cleaves its sole tRNA^LysUUU^ target (29) in a manner dependent on the secondary structure of the anticodon stem-loop as well as the sequence flanking the cleavage site (20). By analogy, MazF-mt1 likely relies on recognition of rRNA structure within the context of the assembled 30S ribosomal subunit.

The 3’ terminus of the 16S rRNA targeted by MazF-mt1 is critical for efficient translation initiation. During canonical translation initiation, the aSD sequence (CCUCCU) near the 16S rRNA 3’ end (Fig. 4) base pairs with complementary AG-rich SD motifs upstream of leadered mRNAs to position the ribosome at the start codon and promote translation initiation. Consistent with the importance of the aSD-SD interaction, its disruption by MazF-mt1 results in severe inhibition of new protein synthesis. Indeed, the phenotype of MazF-mt1 mirrors that of MazF-mt11 previously reported by our laboratory, even though MazF-mt11 cleaves at a different site, C↓ACCU, located 4 nt upstream of the MazF-mt1 cleavage site (Fig. 4). These findings demonstrate that multiple MazF toxins can independently target the 3’ end of 16S rRNA to disrupt translation.

Other toxins appear to target the same region. X-ray structures of two Mtb endoribonuclease toxins, RelE1 (30) and RelS (31), were accompanied by biochemical characterization of their RNA cleavage sites. Although a cleavage consensus sequence could not be derived from these studies, both reported cleavage near the 3’ end of the 16S rRNA: RelE1 at the same site as MazF-mt11 (30) and RelS 1 nt upstream of the MazF-mt1 site (31). Therefore, the 3’ end of the 16S rRNA is a shared target region for select members of the Mtb MazF and Rel toxin families.

Regarding the physiological role of Mtb MazF toxins, several lines of evidence link Mtb MazF toxins to the nonreplicating persistent state characteristic of latent tuberculosis and to the broader Mtb stress response.

First, MazF transcripts are upregulated in drug-tolerant Mtb bacilli derived from human sputum samples (32). Second, MazF transcripts are upregulated upon nutrient starvation (33–35). Third, some MazF toxins are required for drug tolerance, virulence, and stress adaptation in guinea pigs (10). Consistent with a role in stress survival, MazF-mt1 and MazF-mt11 exhibited distinct responses to both the type and duration of exposure to several clinically relevant antibiotics (Fig. 6A-D). The growth arrested state we have documented for both MazF-mt1 (Fig. 2) and MazF-mt11 (22) is also consistent with a nonreplicating persistent state.

We also documented opposing trends for MazF-mt1 and MazF-mt11 expression in a macrophage infection model and under environmental stresses encountered during infection (Fig. 6E,F). Collectively, these differential responses suggest that the two toxins respond to nonoverlapping stresses. The reason for this apparent separation of triggers remains unclear but may provide a mechanism for titrating the magnitude of translation inhibition according to the stress encountered.

Consistent with the canonical role of the SD sequence in translation initiation (reviewed in (36)), we demonstrate that loss of a functional aSD is associated with a dramatic reduction in new protein synthesis. Yet there is a provocative backstory. The transcriptomes of Mtb and *M. smegmatis* contain an unusually high proportion of leaderless mRNAs. Approximately 25% of mRNAs in Mtb and *M. smegmatis*, many of which are polycistronic, are leaderless (35, 37, 38); these transcripts begin directly with the translation start codon and lack an SD sequence. At the individual gene level, ∼14% of annotated transcripts are leaderless in *M. smegmatis* (39) and Mtb (35, 38). Thus, it remains unclear from our metabolic labeling experiment (Fig. 5) whether the small amount of new protein synthesis observed against a background of globally diminished translation represents low level selective translation by ribosomes lacking a functional aSD. However, previous reports serve as a cautionary tale. The proposal that *E. coli* MazF-mediated removal of the aSD sequence generates specialized ribosomes that preferentially translate stress-related transcripts (40, 41) was not supported by subsequent, methodologically rigorous studies (42). Moreover, it is unclear why *M. smegmatis* would contain a similarly high proportion of leaderless transcripts despite lacking an apparent mechanism for generating specialized ribosomes to translate them efficiently. *M. smegmatis* lacks genes encoding Rel family toxins and encodes a single MazF toxin that does not target the 3’ end of the 16S rRNA but instead exclusively cleaves and inactivates tRNA^LysUUU^ (17). Detailed mechanistic studies are therefore warranted to determine whether MazF-mt1 and MazF-mt11 contribute to the nonreplicating persistent state characteristic of latent tuberculosis simply by limiting growth and inhibiting translation, or whether these effects are accompanied by preferential synthesis of a subpopulation of proteins suited to this persistent state.

## Materials and Methods

### Strains, plasmids, and reagents

The *E. coli* strain BW25113Δ6 [*lacI*^q^ *rrnB*_T14_ Δ*lac*-Z_WJ16_ *hsdR*514 Δ*araBAD*_AH33_ Δ*rhaBAD*_LD78_ Δ*chpBIK* Δ*dinJ*-*yafQ* Δ*hipBA* Δ*mazEF* Δ*relBE* Δ*yefM*-*yoeB*] was used for all *E. coli* growth profile and RNA cleavage studies. The *E. coli* strain Mach1-T1 [F^−^ ΔrecA1398 endA1 tonA φ80(lacZ)ΔM15 ΔlacX74 hsdR(r_κ_^−^, m_κ_^+^); Invitrogen] was used for all cloning experiments. The plasmid pBAD33 was used for expression in *E. coli*. All *E. coli* liquid cultures were grown at 37°C in LB medium. Toxin was expressed via an arabinose-inducible promotor in pBAD33-*mazF*-*mt1*. The working concentrations of arabinose and kanamycin were 0.2% and 50 μg/mL, respectively.

The strain mc^2^155 was used for all *M. smegmatis* experiments. For all *M. tuberculosis* experiments we used strain mc^2^6206 (*ΔpanCD ΔleuCD*, generously provided by the William Jacobs Laboratory, Albert Einstein College of Medicine, Bronx, NY) modified by deletion of the *mazEF-mt1* TA module using the ORBIT method (43). The oligonucleotide used to create the deletion was NWO3182, GTG GTC TCC CCG ACG CGG AAC CTG CGA GAG CTA CGA CCA TAA GTC GAG ATG CAG TTT CAA GGT TTG TCT GGT CAA CCA CCG CGG TCT CAG TGG TGT ACG GTA CAA ACC GTC ATC GGA CAG GCT CAC GCT CAA CTT CAC AAC CTA AGA TGC TAC CAG GGT CGT ACC TAG. The plasmid pMC1s was used for expression in both mycobacterial strains. All *M. smegmatis* liquid cultures were grown at 37°C in 7H9 Middlebrook media supplemented with 0.05% Tween 80, 0.5% bovine albumin, 0.2% glucose, and 0.085% NaCl. All *M. tuberculosis* liquid cultures were grown at 37°C in 7H9 Middlebrook media supplemented with 1× oleic acid, albumin, dextrose, catalase (OADC) supplement (BD Biosciences), 0.05% of tyloxapol, 50 µg/mL pantothenic acid, and 100 µg/mL leucine. Toxin was expressed via a tetracycline-inducible promotor in pMC1s-*mazF*-*mt1*. The working concentration of kanamycin was 25 μg/mL.

### RNA isolation for 5’-OH RNA-seq

Total RNA was isolated from *E. coli* freshly transformed with pBAD33-*mazF*-*mt1* and *M. smegmatis* and *M. tuberculosis* transformed with pMC1s-*mazF-mt1* grown to midlogarithmic phase. *E. coli* cultures were grown to an absorbance of 0.3 at 600 nm, after which arabinose was added to a final concentration of 0.2% and the cultures were grown for an additional 2 h at which point cells were harvested. *M. smegmatis* cultures were grown to an absorbance of 0.1 at 600 nm, after which the culture was split into two equal volumes and ATc was added to one culture to a final concentration of 200 ng/ml. The cultures were grown for an additional 5 h. *M. tuberculosis* colonies were split and inoculated into induced and uninduced cultures with the induced receiving ATc to a final concentration of 200 ng/mL. ATc was replenished every 48 h. Cells were harvested 4 and 5 days after induction.

Cells were pelleted at 3000*g* at 4 °C for 10 min, resuspended in TRIzol Reagent (Invitrogen), and lysed in a Precellys Evolution with 0.1 mm silica beads at 10,000 rpm three times for 30 s each time. Lysates were precipitated with ethanol and added to Zymo-Spin IIC columns. After an initial wash with Direct-zol RNA Wash Buffer, genomic DNA was digested with 5 U of DNase I. The RNA was purified following the Direct-zol RNA Miniprep protocol (Zymo Research). To further purify the RNA, RNA was eluted from the column with nuclease-free water, treated with 2 U of TURBO DNase (Invitrogen) for 30 min at 37 °C, and purified using ethanol, a Zymo-Spin IIC column, and the Direct-zol RNA PreWash and Wash buffers as directed by the Direct-zol RNA Miniprep protocol. RNA was eluted from the column with nuclease-free water and the RNA concentrations were measured using a Qubit 4 fluorometer (ThermoFisher).

### MORE RNA-seq and 5’-OH RNA-seq

5’-OH libraries used for MORE RNA-seq and 5’ RNA-seq were constructed as previously described by other members of the laboratory (17, 19–21, 29, 44, 45). The amplified 150-450 bp library fragments were gel purified from a 10% TBE PAGE gel (Invitrogen) and sequenced using the Illumina platform. Using Trim Galore! Version 0.6.10 (46), the resulting FASTQ files had the adaptors sequences and the first 6 nucleotides of the 5’ end trimmed. Additionally, sequences with poor quality scores or less than 20 nts were discarded. Reads were then trimmed to 20 nts using fastx_trimmer from the FASTX Toolkit version 0.0.14 (Hannon Lab, GitHub). The remaining reads were aligned to the corresponding reference genome (NCBI accession HG738867, CP000480, and AL123456 for *E. coli, M. smegmatis*, and Mtb, respectively) using Bowtie version 1.3.1 (47) with a seed length of 1 and no mismatches allowed. Read counts were normalized to sequencing depth and expressed as reads per million of mapped reads. Positions that had 0 counts in the uninduced sample libraries were adjusted to a pseudo-count of 1. Fold change at each position was calculated as the ratio of induced to uninduced read counts. For *E. coli* MORE RNA-seq datasets all hits with a fold change of ≥60 were considered and used to create the WebLogo. For *M. smegmatis* and Mtb datasets, all hits with a fold change ≥25 were considered.

### Labeling of newly synthesized proteins

To assess the effects on new protein synthesis in *M. smegmatis* by the induction of MazF-mt1, triplicate samples were grown to an OD_600_ of 0.1 and split into induced and uninduced samples. ATc was added to induced samples only. After 5 hours and 24 hours of induction cultures were split and azidohomoalanine, AHA, (Sigma) to a final concentration of 50 µM and cultures were incubated for another 3 h. To extract AHA-labelled proteins, cells were pelleted, resuspended in lysis buffer (2% CHAPS and 8 M urea), and lysed using the Precellys Evolution homogenizer using three 30 second cycles at 9000 rpm, interspersed with 2 minute cooling periods. The lysate was centrifuged, and the proteins from the supernatant were linked to an alkyne-containing fluorophore (TAMRA) using the Click-&-Go Protein Reaction Buffer Kit (Vector Laboratories). A total of 5 μg of protein from each sample were resolved on a 17% SDS-PAGE gel, imaged with a Typhoon FLA 9500 (GE Healthcare) image system and quantified using Biorad Image Lab 6.1. After imaging, the gel was silver stained (ThermoFisher) to ensure accurate protein loading.

## Data availability

The RNA sequencing data generated in this study have been deposited in the NCBI Sequence Read Archive under accession number PRJNA1337160.

## Acknowledgements

This work was funded by NIH grant R01AI143760 to N.A.W. and GM135141 T32 grant to T.W.S.

## Author Contributions

The author(s) have made the following declarations about their contributions: Conceived and designed the experiments: T.W.S., U.C., and N.A.W. Performed the experiments: T.W.S., E.A.T., U.C., and T.M. Analyzed the data: T.W.S., E.A.T., U.C., and T.M. Wrote the paper: T.W.S. and N.A.W.

## Supplemental Material

This article contains supplemental material as Supplementary Tables 1, 2, and 3.

